# AAV-mediated neuronal expression of a scFv antibody selective for Aβ oligomers protects synapses and rescues memory in Alzheimer models

**DOI:** 10.1101/2022.06.10.495369

**Authors:** Maria Clara Selles, Juliana T.S. Fortuna, Magali C. Cercato, Luis Eduardo Santos, Luciana Domett, Andre L.B. Bitencourt, Mariane Favero Carraro, Amanda S. Souza, Helena Janickova, Jorge M. de Souza, Soniza Alves-Leon, Vania F. Prado, Marco A. M. Prado, Alberto L. Epstein, Anna Salvetti, Ottavio Arancio, William L. Klein, Adriano Sebollela, Fernanda G. De Felice, Diana A. Jerusalinsky, Sergio T. Ferreira

## Abstract

Brain accumulation of soluble oligomers of the amyloid-β peptide (AβOs) has been implicated in synapse failure and memory impairment in Alzheimer’s disease. Here, we show that treatment with NUsc1, a single-chain variable fragment antibody (scFv) that selectively targets AβOs, prevents the inhibition of long-term potentiation in hippocampal slices and memory impairment induced by AβOs in mice. As a therapeutic approach for intracerebral antibody delivery, we developed an adeno-associated virus vector to drive neuronal expression of NUsc1 (AAV-NUsc1) within the brain. Transduction by AAV-NUsc1 induced NUsc1 expression and secretion in adult human brain slices, and inhibited AβO binding to neurons and AβO-induced loss of dendritic spine loss in primary rat hippocampal cultures. Treatment of mice with AAV-NUsc1 prevented memory impairment induced by AβOs and, importantly, reversed memory deficits in aged APPswe/PS1ΔE9 Alzheimer’s disease model mice. These results support the feasibility of gene-mediated immunotherapy using single-chain antibodies as a potential therapeutic approach in Alzheimer’s disease.

## Introduction

Alzheimer’s disease (AD) is the main form of dementia in the elderly and affects over 35 million people worldwide (1). Remarkable research efforts have been stimulated by the appeal of the amyloid cascade hypothesis, which posits that memory loss in AD results from brain accumulation of aggregates of the amyloid-β peptide (Aβ). However, despite significant progress in our understanding of AD pathogenesis, effective disease-modifying drugs are still lacking (2).

A distinct therapeutic target in AD has emerged in recent years, comprising neurotoxic amyloid-β oligomers (AβOs). The oligomer hypothesis for AD has been proposed as an alternative to the amyloid cascade (3, 4), based on a considerable body of evidence implicating AβOs as causal agents of synapse damage and cognitive impairment in AD (3, 5-6). AβOs accumulate in AD brain and CSF (7-10) and are prominent in various transgenic animal models of AD, including those with little or no amyloid plaque burden (11,12). Experimental exposure to AβOs causes cognitive deficits in animal models and induces major features of AD pathology, including tau hyperphosphorylation, brain inflammation, synapse elimination, and selective nerve cell death (3).

Specific targeting of pathogenic oligomers is expected to provide improved efficacy of Aβ-directed therapies (13). As previously pointed out, immunotherapeutic approaches targeting a broad spectrum of Aβ species (including monomers, oligomers, amyloid fibrils and plaques) may have failed due to a reduction in the effective mass of antibodies that remains available to bind and neutralize the most neurotoxic Aβ species (i.e., oligomers), since a large portion of the antibodies may be lost in off-target interactions (with little or non-toxic monomers, fibrils and plaques) (14).

We here demonstrate the preclinical efficacy of targeting AβOs by NUsc1, an AβO-specific single-chain variable fragment (scFv) antibody selected by phage display from a library of human-derived scFv antibodies. NUsc1 targets a specific subpopulation of AβOs that is highly toxic to synapses, and shows little or no interaction with non-toxic Aβ monomers and with insoluble amyloid fibrils that are constituents of plaques in AD brains (15, 16). An important feature of scFv antibodies is their lack of the Fc domain, which reduces their capacity to activate cellular immune/inflammatory responses (17-19). From a therapeutic standpoint, this is appealing as brain inflammation has been a significant adverse effect in clinical trials of immunoglobulin-based AD immunotherapies (20-23). In addition, the shorter sequence of scFvs compared to intact immunoglobulins facilitates their gene delivery by AAV-based vectors.

Gene-mediated expression of therapeutic antibodies in the brain obviates the need for regimens of repeated injections in patients (18, 19), and allows for local production of antibodies, thus reducing losses associated with inefficient blood brain barrier crossing and systemic clearance mechanisms. AAV vectors are available in various serotypes, are commercially available for use in gene therapy (24) and are currently being tested in numerous clinical trials. For example, Raffi et al. (2014, 2018) have shown that administration of an AAV2 vector harboring the nucleotide sequence for human nerve growth factor induces sustained transgene expression in the brains of control individuals and AD patients without major complications in a two-year follow up period (25, 26). We now describe the development of an AAV9 vector that drives neuronal expression of NUsc1 within the brain and its use to rescue memory in AD models.

## Results

### NUsc1 prevents AβO-induced impairments in hippocampal long-term potentiation and memory

We first tested whether exogenous, recombinant NUsc1 could block the inhibition of hippocampal long-term potentiation (LTP) and memory loss induced by AβOs. We exposed mouse hippocampal slices to 200 nM AβOs and measured LTP responses elicited by high-frequency stimulation at Schaeffer collaterals. Consistent with previous reports (27, 28), LTP was inhibited in hippocampal slices exposed to AβOs. In contrast, slices treated with NUsc1 before application of AβOs exhibited normal LTP (Fig. 1a,b). Control measurements showed that NUsc1 had no effect on LTP in control hippocampal slices (i.e., in the absence of AβOs).

**Figure 1:**
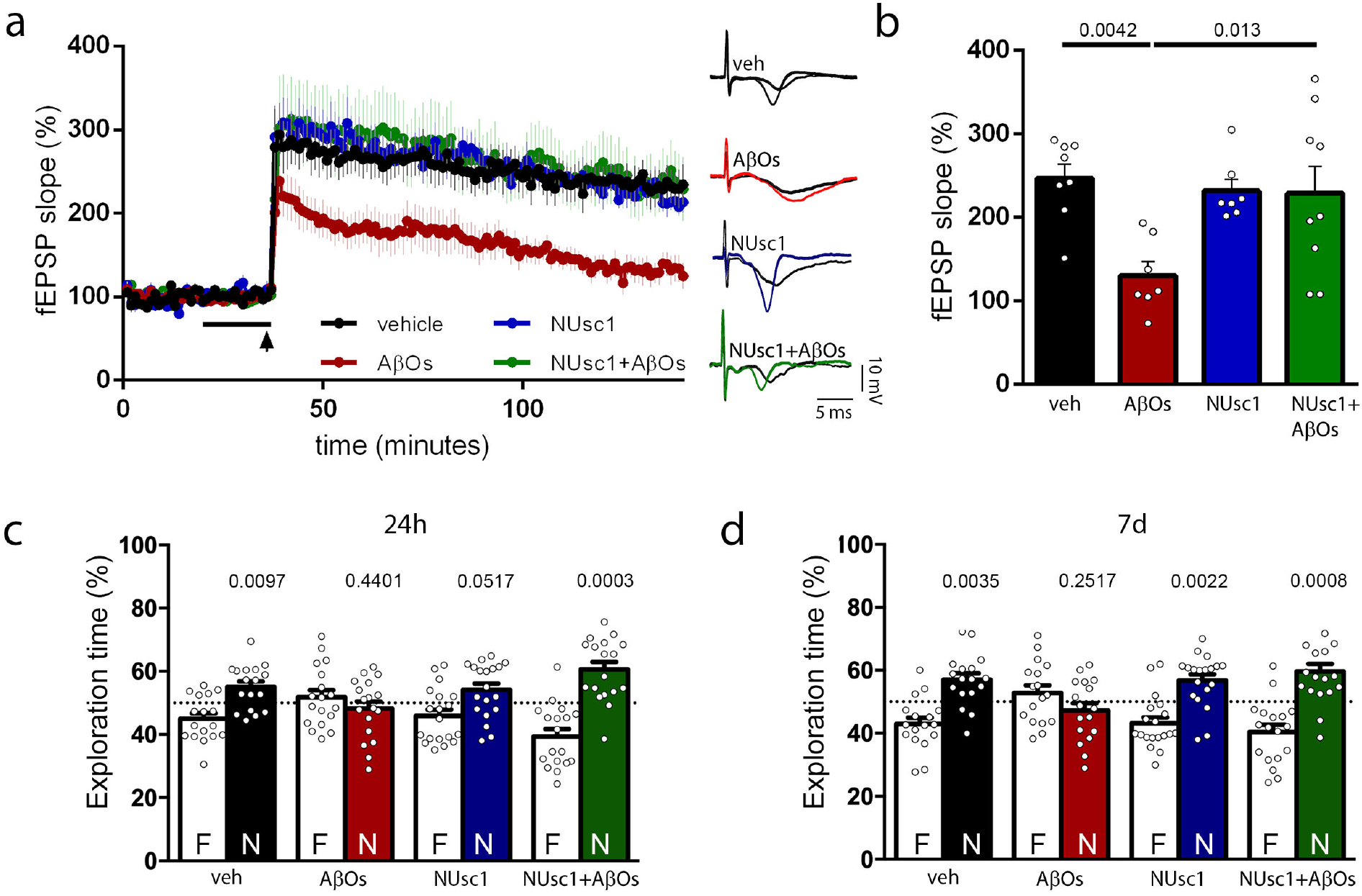
Recombinant NUsc1 prevents AβO-induced inhibition of long-term potentiation (LTP) in hippocampal slices and memory impairment in mice. (a) Long-term potentiation was measured in mouse hippocampal slices. Baseline responses were recorded for 20 minutes, after which slices were perfused for 20 minutes (black horizontal line) with vehicle, purified recombinant NUsc1 (200 pM), AβOs (200 nM), or NUsc1 + AβOs. LTP was elicited by high-frequency stimulation (black arrowhead) at CA3 (Schaeffer collaterals) and recording was performed at CA1 for two hours after stimulus. The main plot shows field excitatory post-synaptic potentials (fEPSP) slopes measured as a function of time in different experimental conditions. Values at each time point are represented as mean ± SE. Representative fEPSP traces before (black lines) and after high-frequency stimulus (colored lines) are illustrated on the right for each experimental condition. (b) Plot of mean fEPSP measured two hours after stimulus. N = 7-9 slices from 5-7 individual mice per experimental condition. Values represent means ± SE, and symbols represent individual slices. P-values for each comparison are shown in the figure; Two-way ANOVA followed by Dunnett‘s post-hoc test. (c-d) NUsc1 prevents AβO-induced memory impairment in mice. Three-month-old Swiss mice received an i.c.v. infusion of NUsc1 (10 fmol) 30 minutes prior to i.c.v. infusion of AβOs (10 pmol). Animals were tested in the Novel Object Recognition (NOR) task 24 hours (c) and 7 days (d) after AβO infusion. Percentages of time spent exploring familiar (F) and novel object (N) are represented by white and colored bars, respectively. Values represent means ± SE and symbols represent individual mice. Color coding is the same in all panels. N= 17-19 total animals per group, tested in 2 independent experiments. P-values for each experimental condition are shown in the graph; two-tailed one-sample Student’s t-test comparing the % of novel object exploration time to the chance value of 50%.

To determine whether NUsc1 could prevent memory impairment induced by AβOs, we treated mice with an intracerebroventricular (i.c.v.) infusion of NUsc1 30 minutes before i.c.v. administration of AβOs and assessed their memory using the Novel Object Recognition (NOR) task. Consistent with our previous reports (28-30), AβO-infused mice exhibited memory impairment both 24 hours and 7 days following AβO infusion (Fig. 1c, d). In contrast, mice treated with NUsc1 exhibited normal performance in the NOR test both 24h and 7 days post-infusion of AβOs.

### Construction of AAV-NUsc1

We next sought to determine whether AAV-mediated transduction and neuronal expression of NUsc1 would be protective in AD models. We constructed an AAV9 vector harboring the nucleotide sequence for NUsc1 (with a single amino acid substitution for expression in eukaryotes; see “Methods”) downstream to a signal peptide (SP) for secretory pathway export and under control of the synapsin I promoter for selective neuronal expression (Fig. 2a). By selectively targeting neurons and employing a single dose of 3×10^9^ vp of AAV-NUsc1, we aimed to prevent overexpression of NUsc1 in the brain, thus minimizing the potential for concentration-dependent antibody self-aggregation, and for induction of aberrant immune or inflammatory responses.

**Figure 2:**
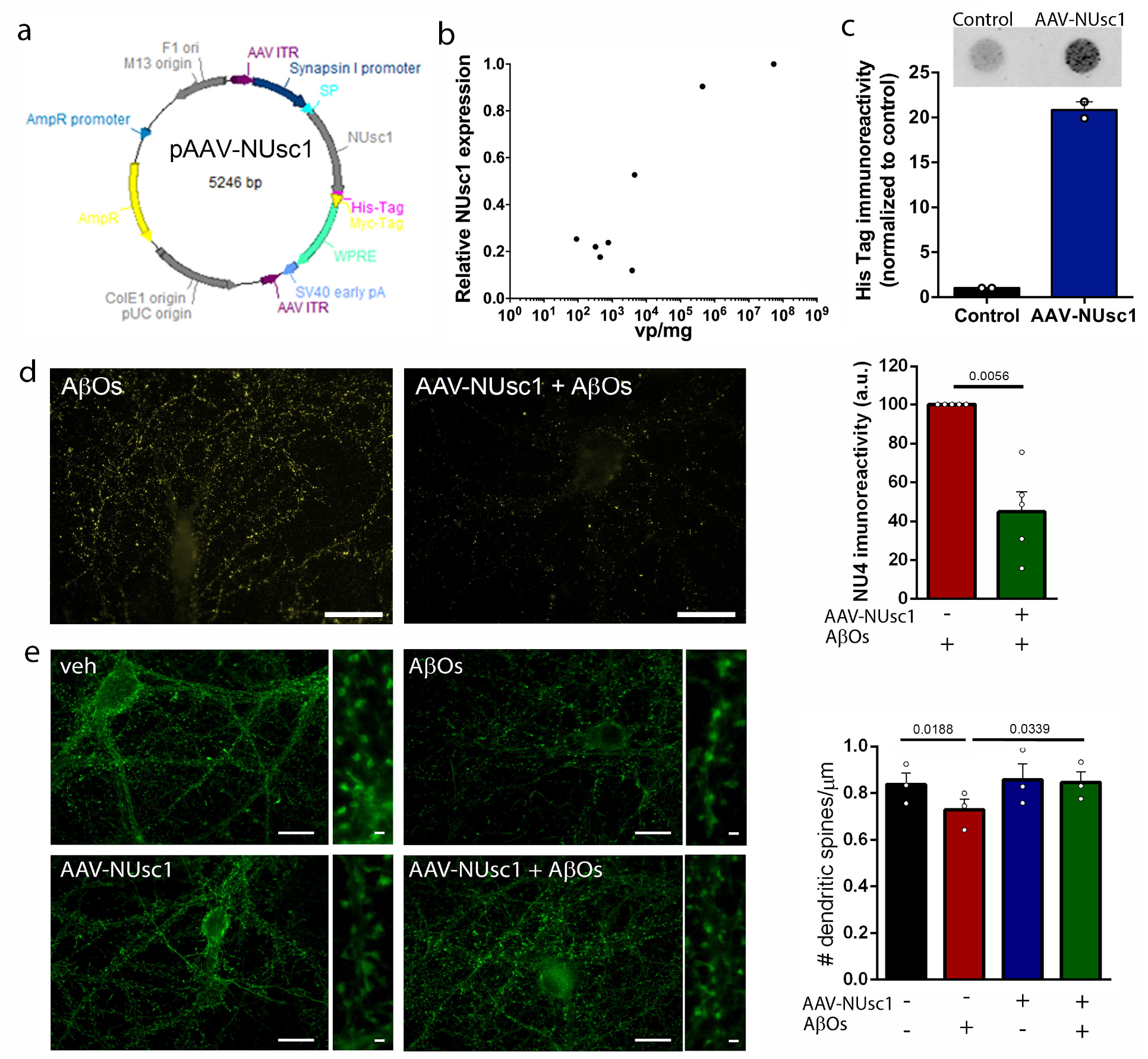
Transduction by AAV-NUsc1 drives NUsc1 expression and secretion in adult human brain slices, decreases AβO binding and prevents dendritic spine loss in rat hippocampal neurons. (a) Map of the pAAV-NUsc1 plasmid used for AAV-NUsc1 vector production. The heavy and light chains of NUsc1 are linked by a flexible linker (VH/linker/VK). NUsc1 has a Signal Peptide (SP) for secretion and two C-terminal (His and Myc) epitope tags. Expression is under control of neuron-specific promoter synapsin I, and the WPRE domain was used to enhance NUsc1 expression. (b) AAV-NUsc1 drives expression of NUsc1 in adult human brain slices in culture. Human cortical slices were infected at 3 days in vitro (DIV) with increasing doses of AAV-NUsc1 vector. At 7 DIV, slices were collected, NUsc1 expression was quantified by qPCR, normalized by 18S ribosomal RNA and plotted as a function of viral particle load per mg tissue. Symbols represent individually treated slices. (c) Dot immunoblot (anti-His antibody) analysis of culture media from vehicle-treated or AAV-NUsc1-infected (10^8^ vp/mg) human brain slices. Representative results (top) and quantification (bottom) of independent cultures from two donors are shown. Values represent means ± SE. (d) Hippocampal cultures were transduced (or not) with AAV-NUsc1 (MOI = 10^4^) at 14 DIV and exposed to AβOs (500 nM) at 20 DIV. AβO binding to neurons was detected using oligomer-specific NU4 monoclonal antibody (40) following 3 hours of exposure to AβOs. Representative images are shown for cultures exposed to AβOs (left) or to AβOs following transduction by AAV-NUsc1 (right). Graph shows integrated fluorescence of bound AβOs (NU4 immunoreactivity, puncta along dendrites). Bars represent means ± SE of 5 experiments (normalized for cultures exposed to AβOs alone) with independent cultures and AβO preparations. Symbols correspond to individual cultures. p-values are shown in the graph; two-tailed paired Student’s t-test. Scale bars correspond to 10 μm. (e) Hippocampal cultures were transduced (or not) with AAV-NUsc1 (MOI = 10^4^) at 14 DIV and exposed to AβOs (500 nM) at 20 DIV. Dendritic spine density was assessed by labeling with Alexa-conjugated phalloidin after 24 hours of exposure to AβOs. Representative images are shown for vehicle-or AβO-exposed cultures previously transduced (or not) by AAV-NUsc1, as indicated in the panels. Scale bars correspond to 15 μm. Insets show optical zoom images of isolated dendrite segments. Scale bars correspond to 1 μm. The number of dendritic spines along 20 μm dendrite segments was quantified. Three dendrite segments from 5 neurons were quantified in triplicate experiments in each experimental condition. Bars represent means ± SE, N = 3 experiments with independent neuronal cultures and AβO preparations; symbols represent independent cultures; p-values for each comparison are shown in the graph; RM one-way ANOVA followed by Dunnett’s multiple comparisons test.

### AAV-NUsc1 induces neuronal expression of NUsc1 in human adult cortical slices

To examine the translational potential of AAV-NUsc1 in AD, we first tested the capacity of AAV-NUsc1 to drive scFv antibody production and secretion in adult human brain tissue (31). Initial tests with an AAV9-mCherry control vector confirmed protein expression in human adult cortical slices in culture (Suppl. Fig. 1). Human cortical slices exposed to increasing titers of AAV-NUsc1 in culture showed dose-dependent expression of NUsc1 (Fig. 2b). NUsc1 protein expression and secretion to the medium were confirmed by dot immunoblot analysis (Fig. 2c). Results indicate that AAV-NUsc1 transduces adult human neurons and drives the expression and secretion of NUsc1.

### AAV-NUsc1 reduces AβO binding to neurons and prevents AβO-induced loss of dendritic spines

To determine whether AAV-NUsc1 could protect neurons from the toxic impact of AβOs, we first transduced mature hippocampal cultures (14 DIV) with AAV-NUsc1. Secretion of NUsc1 to the culture medium was verified by Western blotting at 21 DIV (Suppl. Fig. 2a). Because recombinant NUsc1 has been reported to block AβO binding to neurons (15, 16), we asked whether NUsc1 produced and secreted by AAV-NUsc1-transduced neurons could similarly block neuronal binding of AβOs. Indeed, transduction by AAV-NUsc1 caused ∼ 50% decrease in AβO binding to dendrites in hippocampal neurons (Fig. 2d).

We further investigated whether the reduction in AβO binding could protect neurons from AβO-induced loss of dendritic spines. Neurons exposed to AβOs showed reduced dendritic spine density compared to control cultures, while neurons transduced by AAV-NUsc1 prior to exposure to AβOs exhibited normal dendritic spine density (Fig. 2e).

### AAV-NUsc1 rescues memory in mouse models of AD

As a control experiment prior to assessing the *in vivo* efficacy of AAV-NUsc1, we examined brain distribution and neuronal expression of an mCherry transgene following i.c.v. infusion of an AAV9-mCherry control vector in mice. Results revealed that AAV-mCherry induced expression of mCherry over the entire mouse brain, with prominent expression in the hippocampus and striatum after 8 weeks (Fig. 3a). Having verified that the i.c.v,-infused AAV9 vector successfully transduced neurons in AD-relevant brain areas, we next infused AAV-NUsc1 i.c.v. in mice and verified NUsc1 expression in their brains by qPCR and immunoblotting (Suppl. Fig. 2b,c).

**Figure 3:**
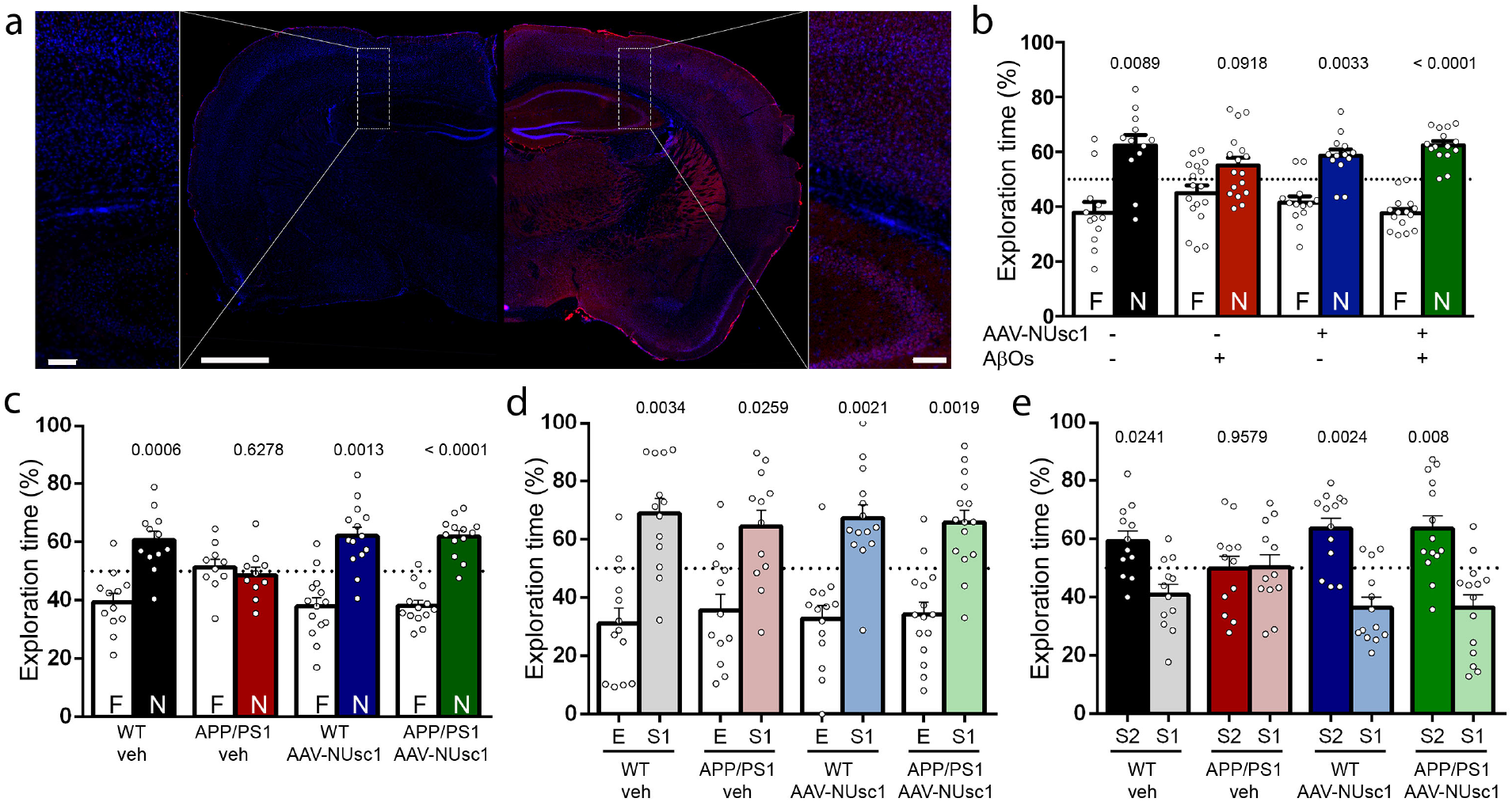
Transduction by AAV-NUsc1 prevents AβO-induced memory loss and reverses memory deficits in aged APPswe/PS1ΔE9 mice. (a) AAV-mCherry (3×10^9^ viral particles) was infused via i.c.v. in 3-month-old Swiss mice. Brain distribution and expression of mCherry were evaluated by immunohistochemistry 8 weeks after infusion. The main image shows a photomontage of coronal sections (−1.82 mm from Bregma) from a control, non-infected mouse (left hemisphere) and an AAV-mCherry-transduced mouse (right hemisphere). Sections were stained with DAPI (blue). Scale bar corresponds to 1 mm. Insets show higher magnification images of areas contained within dashed white rectangles. Scale bar corresponds to 100 μm. (b) Three-month-old Swiss mice received an i.c.v. infusion of 3×10^9^ viral particles of AAV-NUsc1 8 weeks prior to i.c.v. infusion of AβOs (10 pmol). Animals were tested in the NOR task 24 hours after infusion of AβOs. Percentages of time spent exploring the novel object are represented by colored bars. Symbols correspond to individual mice. N= 12-17 animals tested in 3 independent experiments. (c-e) APPswe/PS1ΔE9 mice (9-18 month-old male and female mice) received an i.c.v. infusion of 3×10^9^ AAV-NUsc1 particles. Eight weeks after infection, mice were tested in the NOR task and in the three-chambered social recognition test. (c) Percentages of time spent exploring the novel object in the NOR test are represented by colored bars. Symbols represent data for individual mice. (d) In the three-chambered social recognition test, animals were first habituated in the middle chamber of the apparatus and were then given an option between exploring an empty chamber (E) or a chamber containing a stranger mouse (S1) of the same sex and similar age (white bars labeled “E” versus colored bars labeled “S1” in the graph, respectively). Symbols represent individual mice. (e) In the social novelty part of the task, mice were given the option to explore the already familiar mouse (S1) or a novel mouse (S2). Time spent exploring novel (dark bars) and familiar (light bars, color-coded as in d) mice was quantified. Values represent means ± SE and symbols represent individual mice. N= 10-15 animals tested in 3 independent experiments. P-values for each experimental condition are shown in the graph; two-tailed one-sample Student’s t-test comparing % of exploration time of the novel mouse (S2) to the chance value of 50%.

Two months after i.c.v. infusion of AAV-NUsc1 or AAV-mCherry (used as a control), animals received an i.c.v. infusion of AβOs. When tested in the NOR memory test, both AβO-infused mice (Fig. 3b) and AAV-mCherry-treated/AβO-infused mice (Suppl. Fig. 3) failed the task. In contrast, AβO-infused mice that were transduced by AAV-NUsc1 exhibited normal performance in the NOR task (Fig. 3b). Control experiments showed that transduction by AAV-NUsc1 had no impact on the performance of control, vehicle-infused mice in the NOR test (Fig. 3b).

Finally, we tested the beneficial action of AAV-NUsc1 on memory impairment exhibited by aged APPswe/PS1ΔE9 AD model mice. Memory tests on APPswe/PS1ΔE9 mice (or WT littermates) were performed two months after i.c.v. infusion of AAV-NUsc1. NUsc1 brain levels were evaluated by ELISA in WT and APP/PS1 mice (Suppl. Fig. 2d). APPswe/PS1ΔE9 mice failed both the NOR test and the social memory phase of the three-chambered social recognition test (Fig. 3c,e), but not the sociability phase of the three-chambered test (Fig. 3d). Remarkably, APPswe/PS1ΔE9 mice transduced by AAV-NUsc1 exhibited normal performances in both NOR and social memory tasks (Fig. 3c,e), indicating reversal of memory impairments.

## Discussion

Here, we first found that recombinant NUsc1 prevented the inhibition of LTP in hippocampal slices and memory impairment induced by AβOs in mice. From a mechanistic perspective, AAV-mediated neuronal expression of NUsc1 reduced AβO binding to neurons and blocked AβO-induced loss of dendritic spines in cultured hippocampal neurons. Transduction by AAV-NUsc1 prevented memory deficits in AβO-infused mice and, notably, reversed memory impairments in APPswe/PS1ΔE9 mice. Of relevance from a potential translational perspective, AAV-NUsc1 effectively transduced and drove the expression and secretion of NUsc1 in cultured adult human brain slices.

Gene therapy recently has attracted considerable interest in neurology stemming from its successful introduction in clinical practice for spinal muscular atrophy (32). Moreover, gene therapy has reached clinical trials in a number of neurological disorders, including lysosomal storage diseases, aromatic L-amino acid decarboxylase deficiency disorders and Parkinson’s disease (33). Two gene therapy clinical trials have been registered in ClinicalTrials.gov for AD (NCT00087789 and NCT03634007). While the first approach using an AAV2 vector to induce brain NGF expression failed to prevent cognitive impairment in AD patients (26), it showed no adverse effects. The second approach using an AAVrh.10hAPOE2 to treat APOE4 homozygotes is currently recruiting patients for Phase 1. Preclinical gene therapy studies targeting α-secretase activator/PKC modulator or sAPPα (34) have not yet reached clinical trials.

To our knowledge, this is the first study to utilize a gene therapy approach to achieve sustained neuronal expression of a human-derived scFv antibody that selectively targets a highly toxic subpopulation of Aβ oligomers, and exhibits low affinities for Aβ monomers and fibrils. A number of clinical trials involving Aβ immunotherapies in AD are currently under way, and considerable attention has been given to the recent FDA approval of an antibody targeting multiple Aβ assemblies (Aducanumab) (35). However, considerable controversy remains regarding the benefits of Aducanumab and of other IgGs targeting such heterogeneous Aβ assemblies. Importantly, none of the approaches thus far have taken advantage of the genetic tools available to induce sustained brain expression of an scFv to specifically block the most toxic Aβ species. We found that this strategy led to protection against AβO-induced memory decline in wild-type mice and, significantly, to reversal of age-dependent memory impairment in APPswe/PS1ΔE9 mice. Potential advantages of the current approach include (i) the need for a single infusion of AAV-NUsc1 to achieve sustained brain expression of NUsc1, (ii) local brain expression and secretion of a single-chain antibody devoid of the Fc domain and, thus, less likely to induce aberrant immune/inflammatory responses, (iii) specific targeting of a particular subpopulation of AβOs that is highly toxic to synapses and neurons (15, 16), and (iv) being amenable to further development, including systemic administration of modified AAV vectors capable of effectively crossing the blood-brain barrier (36). Based on our current findings, we propose that development of AAV-NUsc1 represents a step towards reaching immuno-gene therapy for AD.

## Materials and Methods

### Expression and purification of NUsc1

NUsc1 was expressed in the HB2151 E. coli strain with IPTG (Isopropyl b-D-1-thiogalactopyranoside) induction, as described (15). Soluble NUsc1 was purified from both the supernatant and lysate of HB2151 cells using a Protein A affinity column (GE). The solution containing eluted antibody was buffer-exchanged into phosphate-buffered saline (PBS) (Dulbecco’s PBS without calcium or magnesium; Corning), pH 7.4, and concentrated using 10-kDa cutoff centricons (Merck) before storage at -80 °C. The final concentration of purified protein was determined using Bradford reagent (BioRad). Purified NUsc1 was routinely checked by SDS-PAGE and SEC (size-exclusion chromatography).

### AAV-NUsc1 and AAV-mCherry vectors

NUsc1 scFv was subcloned from the original plasmid for expression in bacteria (piT2-NUsc1) by PCR with the following alternative pairs of primers: PCR1: FWD1 (5′-3′) TATTACTCGCGGCCCAGC / REV1 (5′- 3′) CTATGCGGCCCCATTCAG. PCR2: FWD2: (5′-3′) GCTAGCTATTACTCGCGGCCCAGC (with a restriction site for Nhe1) / REV2: (5′-3′) TCGCGACTATGCGGCCCCATTCAG (with a restriction site for Nru1). The PBSKII plasmid was digested with SmaI and used to link the product of each PCR, yielding intermediate plasmids PBSKII-NUsc1 PCR1 and PBSKII-NUsc1 PCR2. The PBSKII-NUsc1 PCR2 plasmid was digested with Nru1, blunt-ended with Klenow polymerase and digested with Nhe1, thus releasing the fragment coding for the open reading frame of NUsc1. This fragment was then cloned into plasmid pA-EAU2 (37) to generate intermediate plasmid pA2-NUsc1, in which NUsc1 is driven by the HCMV promoter. Sequencing of this plasmid revealed a STOP codon between heavy and light chain sequences, which is methylated in bacteria. Since this STOP codon would lead to translation interruption in mammalian cells, site-directed mutagenesis was conducted to replace the STOP codon by a glutamic acid (Glu) codon, resulting in the pA2-NUsc1 plasmid for NUsc1 expression in eukaryotic cells. The NUsc1 protein was then fused to two N-terminal epitope tags (His and Myc tags). In addition, an export signal (SP) was added to promote secretion to the extracellular medium. Finally, the NUsc1 ORF was extracted from pA2-NUsc1 and subcloned into the pA2-SGWA plasmid to obtain plasmid pAAV-NUsc1, in which the NUsc1 ORF is driven by the synapsin I promoter. This plasmid also contains a WPRE sequence to enhance NUsc1 expression (Fig. 2a).

AAV9 vector production was achieved by co-transfection of 293 cells with pAAV-NUsc1, helper plasmid and rep-cap plasmid. Cells were collected 48 h post-transfection and lysed to harvest AAV particles, which were purified by CsCl gradients. The vector was titrated by qPCR. The rAAV particles purified by iodixanol gradients were then quantified; the number of genome-containing particles was assessed by qPCR. Following transduction of mammalian cells, the AAV-NUsc1 vector drives expression of a 30 kDa protein, as expected for the NUsc1 transgene (Suppl. Fig. 2a).

AAV-mCherry control vector was purchased from Virovek (Hayward, CA) and consists in an AAV9 vector harboring the nucleotide sequence for fluorescent protein mCherry under control of the synapsin I promoter.

### Preparation and characterization of AβOs

AβOs were prepared from synthetic Aβ_1–42_ (California Peptide) as previously described (38, 39). Oligomer preparations were routinely characterized by size exclusion HPLC and, occasionally, by Western blots using oligomer-sensitive NU4 monoclonal antibody (40), and comprised a mixture of Aβ dimers, trimers, tetramers, and higher molecular weight oligomers (27, 37, 29). Protein concentration was determined using the BCA assay (Thermo-Pierce).

### Neuronal cultures

Hippocampal cultures were prepared from E18 Wistar rat embryos and were maintained in Neurobasal medium supplemented with B27 (Invitrogen) for 3 weeks as described (41). Cultures were treated with vehicle or AAV-NUsc1 vector using a MOI of 10^4^ at 14 DIV, and were exposed to 500 nM AβOs (or vehicle) at 20 DIV. The supernatant was collected at 21 DIV for determination of the presence of NUsc1 by Western blotting.

### Immunocytochemistry and phalloidin labeling

Cells were fixed and blocked as described (42), incubated with AβO-selective NU4 mouse monoclonal antibody (1 μg/mL; 43) overnight at 4 °C, and incubated for 3 h at 23 °C with Alexa conjugated secondary antibody. Spines were labeled with Alexa-conjugated phalloidin (which binds to spine-localized dense bundles of F-actin) for 20 min at 23 °C, according to manufacturer’s instructions (Invitrogen). Coverslips mounted with Prolong containing DAPI were imaged on a Zeiss Axio Observer Z1 microscope using an EC Plan-Neofluar 63x/1.25 Oil M27 objective. Dendritic spines were quantified manually using NIH Image J.

### Electrophysiological recordings

Electrophysiological recordings were performed as described (44). Briefly, transverse hippocampal slices (400 μm) were prepared and transferred to a recording chamber where they were maintained at 29 °C and perfused with ACSF (2 ml/min flow rate) continuously bubbled with 95% O_2_ and 5% CO_2_. Field extracellular recordings were performed by stimulating the Schaeffer collateral fibers through a bipolar tungsten electrode and recording in CA1 stratum radiatum with a glass pipette filled with ACSF. After evaluation of basal synaptic transmission, a 20 min baseline was recorded every minute at an intensity eliciting a response approximately 35% of the maximum evoked response. Slices were then perfused for 20 min with vehicle, 200 nM AβOs (expressed as Aβ monomer concentration), 200 pM NUsc1, or AβOs+NUsc1. After treatments, LTP was induced by theta-burst stimulation (4 pulses at 100 Hz, with bursts repeated at 5 Hz, and three tetanic 10-burst trains at 15 s intervals). Responses were recorded for 2 h after tetanization and were measured as field excitatory post-synaptic potentials (fEPSP) slopes expressed as percentages of baseline.

### Human cortical slice culture

Cortical tissue was obtained from adult patients submitted to amygdalo-hippocampectomy for the treatment of refractory temporal lobe epilepsy at the University Hospital of the Federal University of Rio de Janeiro. Collection of this tissue for research purposes was under informed consent of the donors and approved by the Institutional IRB of the Federal University of the State of Rio de Janeiro under # CAAE: 69409617.9.0000.5258. A fragment of temporal cortex (surgical access tissue) was collected at the operating room, and was immediately processed and cultured as described (40). Briefly, tissue was sliced at 400 μm using a McIlwain Tissue Chopper. Slices were plated in 24-well plates (1 slice/well) containing 400 μL Neurobasal A (Gibco) supplemented with 1% Glutamax (Gibco), 1% Penicillin/Streptomycin (Gibco), 2% B27 (Gibco) and 0.25 μg/mL Amphotericin B (Gibco) supplemented with 50 ng/mL BDNF (Sigma Aldrich). Cultures were maintained at 37 ºC and 5% CO2.

### Dot immunoblots

Frozen samples of culture media from human cortical slices were thawed and concentrated using 10-kDa cutoff Amicon Ultra-0.5 mL Centrifugal Filters. Samples (20 μg total protein in 200 μL) were spotted onto a nitrocellulose membrane using a vacuum-assisted dot blot apparatus (Bio-Dot Apparatus 1706545, Bio-Rad). Blots were blocked with 5% BSA in Tween-TBS at room temperature for 2 h and incubated at 4 °C overnight with anti-His antibody (1:200; Sigma Aldrich) in blocking buffer. Membranes were then incubated with anti-mouse secondary antibody conjugated to IRDye 800CW (Licor, Lincoln, NE; 1:10,000) at room temperature for 2 h, imaged on an Odyssey Imaging System (Licor) and analyzed using NIH Image J. The integrated density of the dots corresponding to culture media from AAV-NUsc1-treated samples was normalized by the corresponding control.

### Animals and intracerebroventricular (i.c.v.) infusions

Three-month-old male Swiss, or 9-18 month-old APPswe/PS1ΔE9 (or WT C57BL/6 littermates) male and female mice were used. Animals were housed in groups of five per cage with free access to food and water, under a 12 h light/dark cycle with controlled room temperature and humidity. All procedures followed the Principles of Laboratory Animal Care from the National Institutes of Health and were approved by the Institutional Animal Care and Use Committee of the Federal University of Rio de Janeiro (protocol #IBqM 136/15) and the University of Western Ontario (protocol #2016-104 and #2016-103).

For i.c.v. infusion of AAV-NUsc1, AAV-mCherry, AβOs or vehicle, animals were anesthetized for 7 min with 2.5% isoflurane (Cristália, São Paulo, Brazil) using a vaporizer system and were gently restrained only during the injection procedure. A 2.5-mm-long needle was unilaterally inserted 1 mm to the right of the midline point equidistant from each eye and 1 mm posterior to a line drawn through the anterior base of the eyes (29). Swiss mice received 3×10^9^ viral particles of AAV-NUsc1 (in a final volume of 3 μl) 8 weeks before the infusion of 10 pmol AβOs (or an equivalent volume of vehicle). An AAV-mCherry vector was used in control experiments following the same protocol. When indicated, 1.5 μl of purified, recombinant NUsc1 (0.01 pmol) was administered 30 minutes before AβOs via the same i.c.v. injection site. Transgenic APPswe/PS1ΔE9 mice (and WT littermate controls) received 3×10^9^ viral particles of AAV-NUsc1 via i.c.v. two months before behavioral tests. Mice were closely monitored for the duration of the experiment and showed no signs of toxicity following AAV infusion.

### Immunohistochemistry

Animals were anesthetized and perfused with saline, followed by 4% paraformaldehyde in 0.1 M phosphate buffer, pH 7.4. Fixed brains were removed, cryoprotected in increasing concentrations of sucrose, frozen in dry ice and stored at -80 ºC. Coronal sections (40 μm) were obtained on a cryostat (Leica Microsystems) and stored in PBS, pH 7.4. Immunohistochemistry was performed after washing the sections extensively with PBS. Groups of 6 sections per animal were immersed in 0.1% Sudan Black B for 30 minutes, washed three times in PBS and blocked for 2 hours with 0.3% Triton X-100 and 5% BSA in PBS at room temperature. Sections were then incubated overnight with anti-mCherry primary antibody (1:100; ThermoFisher) diluted in PBS, washed and incubated for 2 h with Alexa Fluor 594-conjugated secondary antibody (1:1,000; Life Technologies). After a final washing step, sections were briefly stained with DAPI and mounted with Prolong (ThermoFisher). Images were acquired on a Zeiss Axio Observer Z1 microscope.

### Image processing

Representative images shown in panels 2d, 3a and 1S were processed using Zeiss ZEN software. Contrast was enhanced by adjusting the tonal range of the histograms (i.e. brightness/contrast adjustment) equally across all images being compared. Care was taken to avoid clipping. DAPI was used as a counterstain and thus adjusted more freely, with the goal of not obscuring other channels. Nonetheless, processing of DAPI images was quite similar across all images. No gamma corrections or any other type of non-linear corrections were made in any channel on any of the images shown. All analyses and quantifications were made on raw images prior to any processing.

### Western immunoblots

Forty-eight hours after i.c.v. infusion of AβOs, hippocampi and cortex were dissected and immediately frozen in liquid nitrogen. For total protein extraction, samples were thawed and homogenized in PBS containing a phosphatase and protease inhibitor cocktail (Thermo Scientific Pierce). Protein concentrations were determined using the BCA kit. Samples containing 30 μg protein were resolved in 15% polyacrylamide Tris-glycine gels (Invitrogen) and were electrotransferred to nitrocellulose membranes at 350 mA for 1 h. Blots were incubated with 5% BSA in Tween-TBS at room temperature for 2 h and incubated at 4 °C overnight with anti-His tag primary antibody diluted in blocking buffer. Membranes were then incubated with anti-mouse secondary antibody conjugated to IRDye 800CW (1:10,000) at room temperature for 2 h, imaged using Odyssey® Imaging System (LiCor) and analyzed using NIH Image J.

### RNA extraction and quantitative real-time PCR

Samples were homogenized and RNA was extracted using SV total RNA isolation kit (Promega). RNA purity was determined by the 260/280 nm absorbance ratio. One μg RNA was used for cDNA synthesis using the High-Capacity cDNA Reverse Transcription Kit (Applied Biosystems). qPCR was performed on an Applied Biosystems 7500 RT–PCR system using the Power SYBR kit (Applied Biosystems). Cycle threshold (Ct) values were used to calculate fold changes in gene expression using the 2^-^ΔΔ^Ct^ method (45). The following primers were used to detect the expression of NUsc1: Forward: TCAGCAGAAACCAGGGAAAG; Reverse: CTGCTGATGGTGAGAGTGAAA. Actin-β was used as a housekeeping gene for mice samples and primers used were: Forward: 5’TGTGACGTTGACATCCGTAAA3’; Reverse: 5’GTACTTGCGCTCAGGAGGAG3’. Human slices samples were normalized by 18S ribosomal RNA and primers used were: Forward: ATCCCTGAAAAGTTCCAGCA; Reverse: CCCTCTTGGTGAGGTCAATG.

### NUsc1 ELISA

NUsc1 was quantified by ELISA in hippocampal extracts from WT or APP/PS1 mice. Immediately prior to the assay, aliquots of extracts (prepared in TBS) were diluted in PBS + 2% BSA (Sigma) to a final concentration of 0.6 mg/mL total protein. Samples were added to plate wells coated with rabbit anti-Myc antibody (Sigma; 1:1,000 in PBS) and incubated for 14 h at 4 °C. After washes with PBS + 0.1% Tween-20 (Sigma), mouse anti-His antibody (Sigma; 1:5,000 in PBS + 2% BSA) was added and incubated for 1h. Detection was carried out using anti-mouse IgG-HRP antibody (GE; 1:5000 in PBS + 2% BSA) followed by addition of TMB substrate (Sigma). Reaction was stopped by adding 0.5 M H_2_SO_4_. A NUsc1 standard curve was built using purified recombinant NUsc1 at 0, 1 and 16 nM (R^2^ =0.99) (15). Optical density values from control (vehicle-infused) mice were averaged and the resulting mean value was subtracted from signals obtained from AAV-NUsc1 treated mice to obtain the mass of NUsc1 in each extract.

### Novel object recognition task

The task was performed in an open field arena measuring 30 × 30 × 45 cm (W x L x H). The floor of the arena was divided by lines into nine equal rectangles. Test objects were made of glass or plastic and had different shapes, colors, sizes, and textures. During sessions, objects were fixed to the box to prevent displacement caused by exploratory activity of the animals. Previous tests showed that none of the objects used evoked an innate preference. Before training, each animal was submitted to a 5 min habituation session to freely explore the empty arena. Training consisted of a 5-min session during which animals were placed at the center of the arena in the presence of two identical objects. The amount of time spent exploring each object was recorded. Sniffing and touching the object were considered exploratory behavior. The arena and objects were cleaned thoroughly between trials with 40% ethanol to eliminate olfactory cues. In the test session, performed two hours after training, one of the two objects used in the training session was replaced by a new one. Time spent exploring familiar and novel objects was measured. Results were expressed as percentage of time exploring each object during the test session and were analyzed using a one-sample Student’s t-test, comparing the mean exploration time for each object against the fixed (chance) value of 50%. An animal that recognizes the familiar object (i.e., that learns the task) explores the novel object > 50% of the total time.

### Three-chambered social recognition task

Each animal was positioned in an apparatus divided into three equally sized chambers and tested along three sessions lasting 5 min each. The first session consisted in free exploration of the middle chamber. The second session evaluated social interaction by quantifying the time spent exploring each of the side chambers – one containing a small empty wire cage and the other containing an identical cage containing another mouse of the same sex and similar age as the test animal inside. Finally, the third session consisted in placing a novel mouse in the empty cage on the opposite chamber and quantifying the amount of time spent exploring the familiar and novel mice, to evaluate social memory.

### Statistical analysis

Results were tested using the D’Agostino & Pearson omnibus normality test and are represented as means ± SE. All analyses were performed using GraphPad Prizm software. Outliers were identified and were removed when applicable. Sample sizes and statistical tests used to analyze the results are specified in the corresponding Figure Legends. Detailed information on statistical analysis is presented in Supplementary Table 1.

## Acknowledgements

S.T.F. and F.G.D.F. were supported by grants from National Council for Scientific and Technological Development (CNPq/Brazil), Fundação de Amparo à Pesquisa do Estado do Rio de Janeiro (FAPERJ/Brazil) and National Institute for Translational Neuroscience (Brazil). S.T.F. and D.A.J. were jointly supported by a binational research grant from CNPq/CONICET. ASe was supported by the Sao Paulo Research Foundation (FAPESP) grant #2014/25681-3. M.A.M.P and V.F.P. received support from the Alzheimer’s Society of Canada and the Canadian Institutes of Health Research (CIHR, PJT 162431 and PJT 159781). ALBB received a pre-doctoral fellowship from CNPq. M.C.S. received pre-doctoral fellowships from CNPq and FAPERJ, and travel grants from International Society for Neurochemistry, Company of Biologists, and Coordenação de Aperfeiçoamento de Pessoal de Nível Superior. We thank Drs. Moses Chao and Mauricio M. Oliveira for critical reading of the manuscript and discussions.

## Author Contributions

M.C.S., D.A.J. and S.T.F. designed the study. A.Se., W.L.K., S.T.F. developed the recombinant NUsc1 scFv. D.A.J., A.L.E., A.Sa. and S.T.F. conceived constructs and vector development. M.C.S., J.T.S.F., M.C.C., L.E.S., L.D., A.L.B.B., M.F.C., A.S.S. and H.J. performed research. M.C.S., J.T.S.F., M.C.C., L.E.S., L.D., A.L.B.B., O.A., F.G.D.F. and S.T.F analyzed data. J.M.S., S.A.-L., V.F.P., M.A.M. P., A.L.E., A.Sa., A.Se. and W.L.K. contributed reagents, materials and analysis tools. M.C.S., J.T.S.F., L.E.S., A.L.E., A.Sa., A.Se., O.A., W.L.K., F.G.D.F., D.A.J. and S.T.F. analyzed and discussed the results. M.C.S., W.L.K. and S.T.F. wrote the manuscript with contributions from other authors.

