## Supplementary Figures for "AAV-mediated neuronal expression of a scFv antibody selective for Aβ oligomers protects synapses and rescues memory in Alzheimer models"

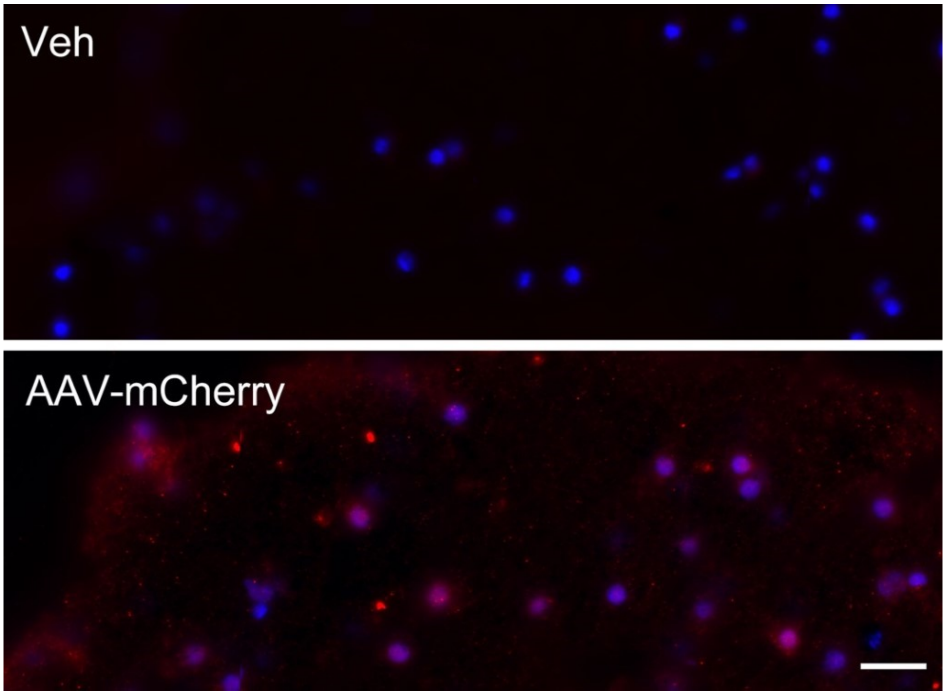


Supplementary Figure 1: Human adult cortical slices (see “Methods”) were infected with AAV-mCherry ($10^8$ vp/mg). mCherry fluorescence was detected by immunohistochemistry. Vehicle-treated control (upper panel) or AAV-mCherry-transduced (lower panel) human slices were imaged at 40x. Scale bar corresponds to 20 μm.

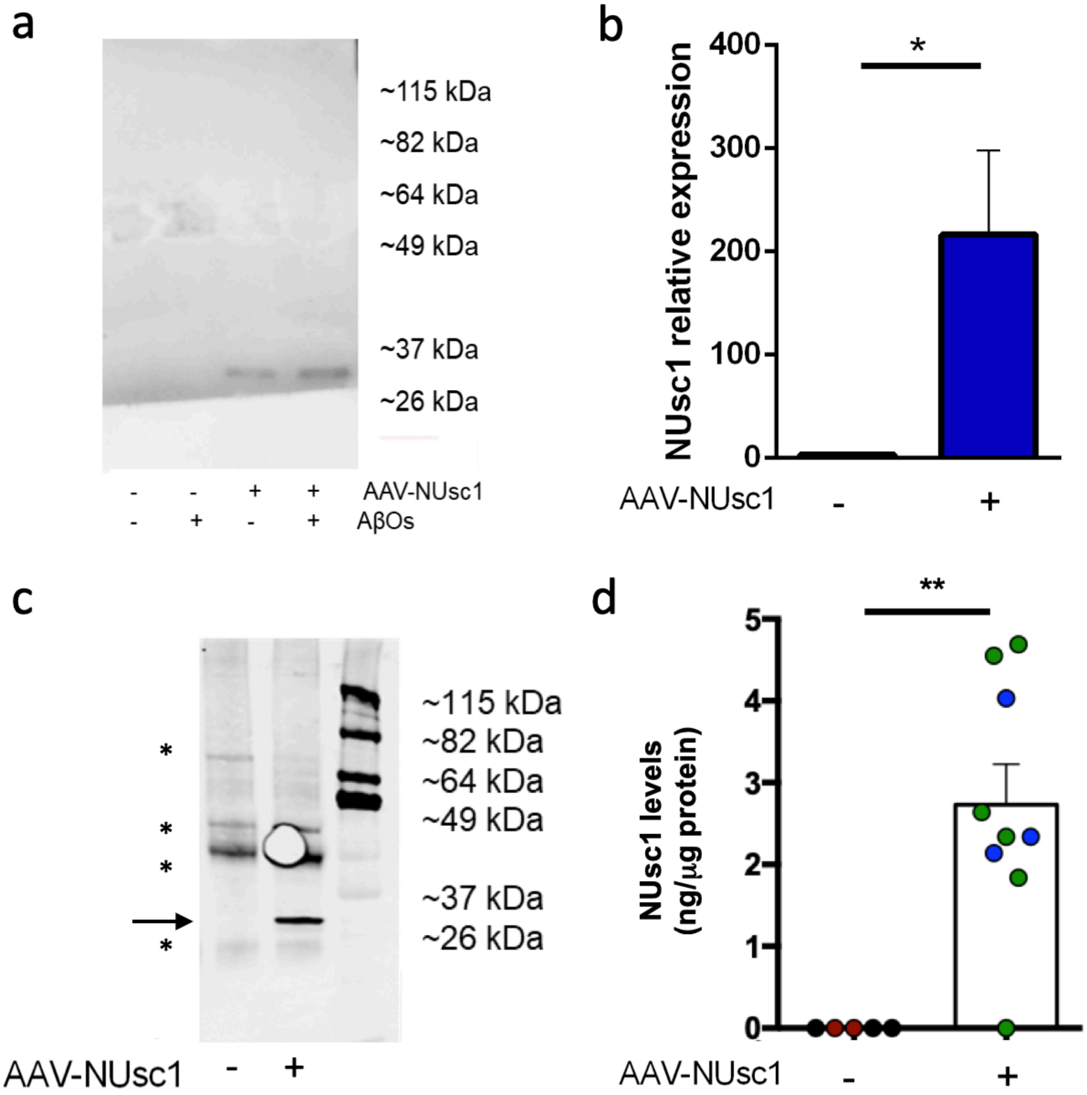


Supplementary Figure 2: (a) Western blot of culture media from rat hippocampal neuronal cultures transduced using a MOI of $10^4$ AAV-NUsc1 vp/cell. NUsc1 (~30 kDa) was detected with anti-His antibody. (b-d) NUsc1 expression in the brains of mice that received an i.c.v. infusion of $3x10^9$ vp of AAV-NUsc1, determined by (b) qPCR (means ± SE. N = 13 animals per group; two-tailed Mann-Whitney test), (c) Western blot (arrow points to band corresponding to NUsc1; asterisks mark nonspecific bands), and (d) sandwich ELISA employing 6xHis and Myc tags present in NUsc1 (black symbols: vehicle-treated WT mice; red: vehicle-treated APP/PS1 mice; blue: AAV-NUsc1-treated WT mice; green: AAV-NUsc1-treated APP/PS1 mice. Student's t test. * $p<0.05$, ** $p<0.01$

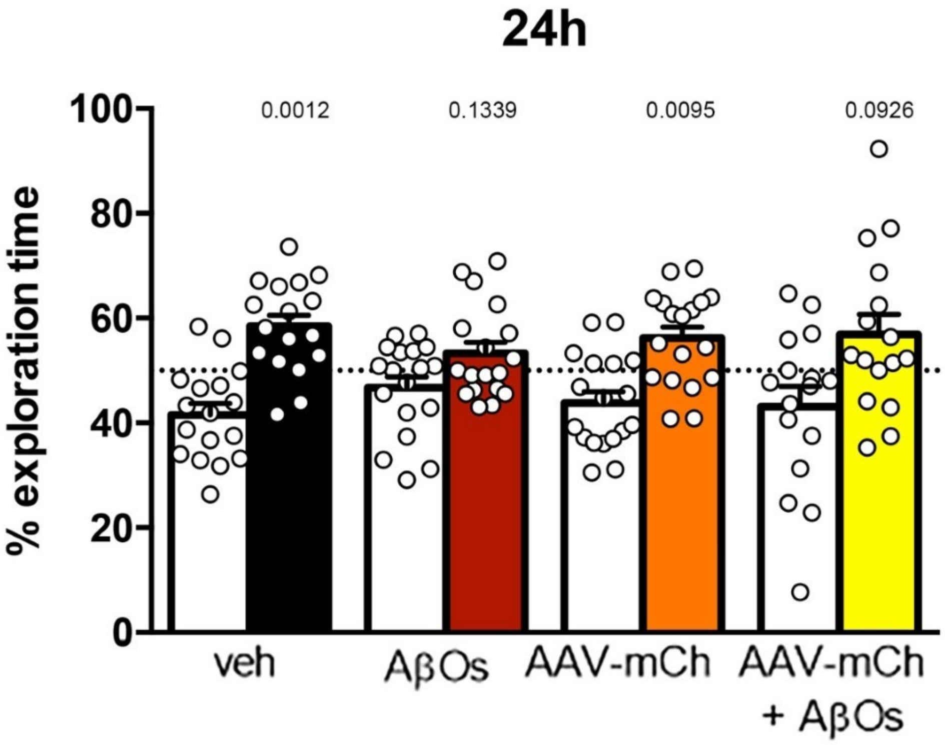


Supplementary Figure 3: Transduction by AAV-mCherry does not protect mice from AβO-mediated memory loss. Three-month-old Swiss mice received an i.c.v. infusion of $3x10^9$ viral particles of AAV-mCherry 8 weeks prior to i.c.v. infusion of AβOs (10 pmol). Animals were tested in the NOR task 24 hours after infusion of AβOs. The percentages of time spent exploring the novel object are represented by colored bars. Values represent means ± SE. N = 16-18 animals tested in 2 independent experiments. Two-tailed one-sample Student’s t-test comparing % of time spent exploring the novel object to the chance value of 50%.
